# Atypical p38 Kinase Signaling in Retinal Vascular Damage and Recovery

**DOI:** 10.1101/2025.08.29.672721

**Authors:** Lillian Schulz, Abby E. Young, Fang Liu, Sanjana Gattineni, Sneha S Ghosh, Eugene Douglass, Heike Kroeger, S. Priya Narayanan, Neil J. Grimsey

## Abstract

Despite the life-changing impact of anti-VEGF therapies, vascular retinopathies continue to affect millions of patients worldwide. To better understand disease progression, it is essential to define alternative mechanisms that contribute to retinal vascular dysregulation. Mitogen-activated protein kinase (MAPK) p38 plays a significant role in regulating vascular homeostasis, angiogenesis, and retinal disease progression, yet effective therapeutic targeting remains elusive. An alternative atypical p38 signaling pathway is mediated by interaction with the adaptor protein TGF-beta activated kinase 1, binding protein 1 (TAB1). Atypical p38 can be activated by ischemia or by inflammatory G protein-coupled receptors (GPCRs) to regulate inflammation and vascular homeostasis. However, atypical signaling has not been investigated in the context of vascular retinopathies, specifically oxygen-induced retinopathy (OIR). Here, we utilized a genetic knock-in mouse to block atypical p38 activity (Tab1^KI^) to explore retinal damage during OIR. We report that Tab1^KI^ mice display significantly reduced vaso-obliteration and limited neovascularization relative to wild-type C57BL6 controls. Retinal RNAseq analysis revealed distinct transcriptional regulation in the Tab1^KI^ mouse. Suppression of atypical p38 reduced Mef2c signaling, which in turn enhanced microglial activation and inflammation. Despite the upregulation of proangiogenic markers, the increase in endothelial markers does not correlate with the dysregulated growth of blood vessels. Instead, Mef2c suppression reduces pathological angiogenesis, while increasing physiological vascular regrowth. The combined data suggest that the selective inhibition of atypical p38 signaling could block the enhanced pathogenic neovascular tuft formation without impacting blood vessel repair. Further investigation will clarify how atypical p38 controls neovascular responses, potentially revealing new treatment strategies for vascular retinopathies.

**NEW AND NOTEWORTHY:** MAPK p38 drives ocular injury and inflammation, but the contribution of Tab1-dependent atypical p38 in retinal injury has not been investigated. Using a mouse genetic knock-in model to block atypical p38 during oxygen-induced retinal injury, we show that atypical p38 enhances vascular damage and neovascular tufting. Blockade of atypical p38 protects mice from retinal vascular damage.

## INTRODUCTION

Pathologic ocular neovascularization is driven by ischemic signaling in diabetic retinopathy (DR), neovascular age-related macular degeneration (nAMD), and retinopathy of prematurity (ROP). Ischemic retinal damage is the leading cause of blindness in adults (proliferative DR and nAMD) and neonates (ROP). In DR and ROP, retinal damage occurs in two phases. First, non-proliferative vascular remodeling leads to vascular damage and loss of blood vessels (vaso-obliteration), which induces a hypoxic environment. The subsequent ischemic phase triggers dysregulated vascular repair, mediated by increased inflammatory signaling and proangiogenic growth factors. Pathological retinal neovascularization leads to weakened blood vessels and aberrant, hyperproliferative endothelial growth, which damages the retina and contributes to vision loss in millions of patients.

Vascular dysregulation is primarily driven by vascular endothelial growth factor (VEGF) signaling [1-4]. Anti-VEGF therapies remain the cornerstone for suppressing aberrant angiogenesis, vascular tufting, and macular edema [2, 5, 6]. However, anti-VEGF therapies do not promote vascular repair, and a substantial group of patients exhibit inadequate, transient responses or develop resistance to anti-VEGF therapies [7-9]. This highlights the importance of identifying the cellular effectors that contribute to retinovascular damage and vision loss. One such effector is the p38 mitogen-activated protein kinase (MAPK) family, which occupies a central node in cellular stress and inflammatory responses [10, 11]. In the eye, p38α activation has been implicated in endothelial dysfunction, glial activation, and pro-inflammatory signaling across DR [12-21], glaucoma [22-24], AMD [25, 26], and ischemic retinopathies such as ROP [27, 28].

Canonical p38 activation is induced via a three-tiered cascade in which upstream kinases MKK3/6 phosphorylate p38α at Thr180/Tyr182 [10]. However, emerging evidence describes a distinct MKK-independent “atypical” activation pathway mediated by TAK1-binding protein 1 (TAB1) [10]. TAB1 docks directly onto two discrete domains of p38α, inducing its autophosphorylation and selectively propagating pathological signaling[29]. The TAB1 interaction is specific to p38α, and Tab1 cannot bind the other isoforms p38β, γ, or δ [30]. Our previous studies characterized a novel mechanism in which inflammatory vascular G protein-coupled receptors (GPCRs) in the regulation of vascular homeostasis and inflammatory signaling [31, 32]. Despite the evidence for TAB1-dependent signaling in vascular damage and repair and its critical role in hypoxic signaling during ischemia-reperfusion injury [29, 33], the contribution of atypical p38 in retinal injury has not been explicitly examined.

To dissect the role of atypical p38 signaling in retinal neovascular diseases, we employed a Tab1 knock-in (Tab1^KI^) mouse harboring point mutations that abrogate Tab1–p38α binding while preserving canonical MKK3/6-driven activation [29]. We subjected wild-type and Tab1^KI^ mice to the oxygen-induced retinopathy (OIR) model - a biphasic paradigm of hyperoxia-induced vaso-obliteration followed by hypoxia-driven, pathological neovascularization - to test the hypothesis that Tab1-dependent p38 autophosphorylation exacerbates vascular injury. Our findings demonstrate that selective loss of atypical p38 activation confers significant protection against vascular obliteration and neovascular tufting in OIR, despite intact canonical p38 signaling and preserved hypoxic and angiogenic responses. Transcriptomic profiling further revealed that disruption of the Tab1–p38α axis diminishes a discrete subset of p38 target genes implicated in inflammation and angiogenesis, identifying atypical p38 signaling as a novel molecular driver of retinovascular pathology and vascular repair.

## Methods and Materials

### Reagents and Antibodies

The following antibodies were used: Rabbit anti-GFAP (DAKO, # Z0334), Mouse anti-CD31 (DAKO, #M0823), Rabbit anti-glutamate synthase (Abcam, #ab228590), Rabbit anti-Iba1 (WAKO, #019–19741), Isolectin-B4 Alexa-Fluor 488 (Invitrogen, #I21411), Mouse anti VEGFA (Santa Cruz, #SC-7269) Rabbit total p38 (Cell Signaling Technologies, #9212S), Rabbit phospho-p38 (Cell Signaling Technologies, #4511S), Mouse anti-Tab1(Gentex, #GTX107571), Rabbit anti-phospho-MKK3/6 (Cell Signaling Technologies, #9231S), rabbit anti-total-MKK6 (Cell Signaling Technologies, #8550S). Donkey anti-Rabbit IgG Alexa Fluor 488 (Invitrogen, # A-20216), Donkey anti-Mouse IgG (H+L) Alexa Fluor 555 (Invitrogen, #A-31570), Goat anti-rabbit IgG (H+L)-HRP Conjugate (Bio-Rad Laboratories, 1706515), Goat anti-mouse IgG (H+L)-HRP Conjugate (Bio-Rad Laboratories, #1706516). PhosTag (Wako Pure Chemical Industries, #AAL-107).

### Mouse Strains

The wt C57BL/6 female mice were purchased from Charles River Laboratory and used to reconstitute the Tab1^KI^ (V390A, Y392A, V408G, and M409A) mice originally provided by the lab of Dr. Michael S. Marber (KCL, UK). Heterozygous KI mice were back-crossed with WT mice for 4 generations and genotyped to confirm Tab1-mutations using the primers: Neo_Del_F: 5′-GCTGGCCTTGCTCAACTCCAG-3′, Neo_Del_R: 5′-GACCATCTGTCTCATACCTGACCTCAC-3′ Genotyping using a mouse direct PCR kit, as per the manufacturer’s instructions (Sellechem, #B40015). Homozygous wt or KI mice were used for all experiments. All animal experiments were performed according to the National Institutes of Health Guide for Care and Use of Laboratory Animals and were approved by the University of Georgia Institutional Animal Care and Use Committee (IACUC).

All animal procedures were approved by the Institutional Animal Care and Use Committee (IACUC) at Charlie Norwood Veteran Affairs Medical Center (CNVAMC), Augusta, GA, Augusta University, Augusta, GA, and University of Georgia, Athens (UGA), GA. In this study, all efforts were made to minimize animal suffering. Wild-type and Tab1^KI^ C57BL/6 mice were maintained in the animal facility at CNVAMC and UGA. The experimental model of OIR was induced according to the methods standardized in our laboratory[34]. Briefly, on postnatal day 7 (pn7) newborn mice and their nursing mothers were placed in a 70% oxygen chamber (hyperoxia phase) for five days. On pn12, mice were returned to room air and maintained until pn17 (hypoxic phase). Mice were sacrificed, and ocular globes or retinal tissues were collected and analyzed as described below.

### Immunoblotting

pn17 RA, and pn17 OIR were euthanized by ketamine/xylazine injection before retina dissection in situ and immediate freezing in liquid nitrogen. Note, the right eye was kept for RNAseq analysis, and the left eye was processed for immunoblotting as follows. Retinae were thawed and lysed directly in Triton X-100 lysis buffer (1% Triton X-100, 50 mM Tris-HCl pH 7.0, 100 mM NaCl, 5 mM EDTA, 10 mM EGTA, 10 mM β-glycerophosphate, 10 mM NaPP, 200 µM mM NaVO_4_, 1 μg/ml leupeptin, 1 μg/ml aprotinin, 1 μg/ml trypsin protease inhibitor, 1 μg/ml pepstatin, 10 μg/ml benzinamide), quantified by BCA assay as per manufacturers instructions, normalized and mixed with 1X Laemmli sample buffer plus 100mM DTT. Proteins were resolved by SDS-PAGE and processed for immunoblotting. Immunoblots were imaged on a Bio-Rad ChemiDoc Imaging System, and quantified using Fiji (National Institutes of Health, Bethesda, MD).

### Immunofluorescence of Retinal Flat Mounts

Flatmounts were prepared as previously described[35, 36]. Briefly, pn17 RA, and pn17 OIR were euthanized by ketamine/xylazine injection, ocular globes were enucleated and fixed in 4% PFA before retinal dissection[37]. Flat mounts were permeabilized using 0.3% Triton-X 100 in PBS and blocked in 0.3% Triton-X 100, 0.2% BSA and 5% goat serum in PBS, before O/N Isolectin B4 staining and mounting onto glass slides using ProLong Gold^TM^. Slides were imaged using a Zeiss LSM800 confocal microscope.

### Retinal Vascular Analysis

Sholl analysis was performed using Fiji software as previously described [35, 38]. Briefly, retinal flat mount images were converted into binary images, masked, and a radius encompassing all vasculature (r∼1000) was established. The neuroanatomy plug-in was used to run a Sholl analysis program. Data was compiled into Microsoft Excel software and integers across repeated radii were averaged to generate a graphical representation (OIR n=6, RA n=4).

### Retinal Sections (staining and immunohistochemistry)

Enucleated ocular globes processed for sectioning as previously described. Briefly, tissues were fixed in PFA and cryoprotected in OCT media (optimal cutting temperature). Retina cryostat sections (10µm thickness) were permeabilized in Triton -X 100 (1% in PBS) for 10 min and blocked in 10% normal goat serum containing 1% BSA for 1 h. Sections were then incubated overnight in primary antibodies at 4 °C. The next day, the sections were incubated at room temperature for 1 h in fluorescent conjugated secondary antibodies and mounted with a DAPI-containing medium (ProLong Gold^TM^). Alternatively, ocular globes were fixed in Bouin’s solution (Millipore, # HT10132) before embedding in paraffin and sectioning. Hematoxylin (Agilent, # S330930-2) and Eosin (Agilent, # CS70130-2) staining as pr the manufacturer’s instructions.

### Tuft Counting

Ganglion cell layer tufts were hand-counted from images of H&E-stained OIR retinal sections. 19 wild-type and 18 Tab1^KI^ retinal sections were counted, n = 6 wt and 6 KI. Data was analyzed in GraphPad Prism 7.0 software for outliers and significance. No outliers were identified. Significance was evaluated using an unpaired t-test.

### Microglia Counting

Ramified and amoeboid microglia were hand-counted from immunofluorescence images of OIR retinal sections, using previously established reference images[39]. 18 images from wild-type mice and 36 images from Tab1^KI^ mice were assessed, giving n-values of 18 and 36, respectively, from 18 mice per group.

### Tortuosity

Analysis was performed in Fiji software by measuring true vessel lengths and the direct distance between start and end points. Seven veins each from four wild-type and four Tab1^KI^ room air flat mounts were measured, giving a room air n of 28. Seven veins each from six wt OIR flat mounts and eight Tab1^KI^ OIR flat mounts were measured, providing n values of 42 and 56, respectively. Direct paths were divided by the corresponding length. Vessel Maps were created by generating skeletonized images of vascular architecture in Fiji software and importing into Adobe Photoshop (Adobe Inc., San Jose, California) to edit for clarity.

### RNA Seq sample preparation

Retinae were collected as described above, thawed directly into TRIzol, and mRNA isolated using Qiagen mRNA isolation kit, as per kit instructions (RNeasy universal kit, Qiagen, #74106). Sample mRNA was sequenced by NGS sequencing (Novogene), using the Mus Musculus(GRCm39/mm39) reference sequence.

### Computational Methods: Sample Processing, Dimensionality Reduction, and Differential Expression Analysis

RNA-seq was performed on mouse retinal samples derived from four experimental groups: wt RA, Tab1^KI^ RA, wt OIR, and Tab1^KI^ OIR, four retinas per group (the other retina from each mouse was used for immunoblots as described). Log-transformed reads per million (log10[RPM + 1]) were used for downstream analyses, with gene-wise z-scores computed for clustering and visualization. Dimensionality reduction was performed using UMAP on the transposed z-score matrix, enabling the projection of sample relationships into two dimensions. Samples were grouped based on genotype and treatment condition, and interactive UMAP plots were generated using the plotly package, while static scatter plots with sample labels provided clarity for print figures. For differential expression, we implemented both a custom t-test–based loop and a DESeq2-based pipeline. The custom loop calculated log fold change and log-transformed p-values for each gene by comparing mean expression between groups (e.g., Tab1KI_OIR vs WT_OIR), while the DESeq2 approach involved back-transforming log-RPM values to raw count estimates, applying integer rounding, filtering out genes with excessive zero counts, and estimating dispersion and fold changes under a negative binomial model. False discovery rates were computed using both Benjamini-Hochberg correction (Q-value) and local FDR via the fdrtool package. Both statistics were calculated after removing NA p-values, with local false discovery rates inserted back into the results table for consistency across genes. Significant genes were visualized with volcano plots, and top candidates were annotated, particularly those corresponding to curated lists of canonical and atypical P38 signaling targets.

### Pathway Enrichment, Hallmark Signatures, and Transcription Factor Activity

To explore pathway-level responses, we loaded GSVA-derived Hallmark pathway enrichment scores for all samples and computed group-level averages. Heatmaps were generated for the top 10 pathways showing the largest intergroup differences, with particular focus on P38-relevant transcriptional programs/gene-sets such as IL6_JAK_STAT3_SIGNALING, TNFA_SIGNALING_VIA_NFKB, TGF_BETA_SIGNALING, HYPOXIA, and PROTEIN_SECRETION. Pathway gene sets were retrieved from the MSigDB database via the msigdbr package for *Mus musculus* and filtered to include only genes expressed in the dataset. For select Hallmark pathways and retina-specific gene sets, z-scored expression matrices were exported to CSV files, enabling independent visualization (e.g. violin plots).

For each Hallmark pathway of interest, gene-level expression was extracted and visualized across conditions using z-score heatmaps, prioritizing genes with high inter-sample variance. A similar strategy was applied to retina-specific gene sets curated from Human Phenotype Ontology terms (e.g., RETINAL_DETACHMENT, CHORIORETINAL_ATROPHY). Transcription factor (TF) activity was inferred using the VIPER algorithm with DoRothEA transcriptional networks applied to RNA-seq expression data[40, 41], resulting in regulon activity scores for each sample. A panel of TFs downstream of P38 kinase signaling according to the KEGG database (e.g., MEF2A/C/D, STAT1/3/6, NFATC4, TP53, CEBPB) was evaluated, with differential activity visualized using both heatmaps and volcano plots. The TF panel included both canonical (e.g. MEF2 family, STATs) and noncanonical (NFATC4, GATA4) effectors, with regulon activity summarized by averaging across biological replicates prior to plotting. Expression and activity patterns for these TFs were compared across all conditions to identify regulators perturbed by Tab1^KI^ status and OIR exposure.

### Network Analysis, Functional Enrichment, and Effector Gene Programs

STRING-based network analysis was used to explore protein-protein interactions among top differentially expressed genes. Gene lists were filtered to retain those with log fold change exceeding ±0.4 and mapped to STRING identifiers using the STRINGdb package (v11.5, species = *Mus musculus*). Networks were pruned to retain high-confidence interactions (combined score >150), and disconnected nodes were removed. The resulting graphs were visualized using ggraph, and node degree was used to identify central hub genes. These top-ranked genes were converted to Entrez IDs using org.Mm.eg.db and submitted for Gene Ontology (GO) enrichment analysis using the clusterProfiler package. Functional terms were summarized via dot plots to highlight enriched biological processes among key regulatory genes. Finally, curated gene sets representing angiogenesis (e.g., Vegfa, Angpt2, Pdgfb, Flt1) and immune signaling (e.g., Il6, Tnf, Ifng, Csf1– 3) were extracted and visualized across experimental conditions using heatmaps of z-scored expression. These analyses provided insight into how atypical P38 signaling may alter effector programs governing cytokine secretion, vascular remodeling, and transcriptional regulation in the retina. All analyses were conducted in R using standard and custom-built functions, with key packages including umap, DESeq2, viper, gplots, plotly, msigdbr, STRINGdb, clusterProfiler, fdrtool, and enrichplot. Retinal cell type gene lists derived from Zarkada *et al* 2021[42].

### Phos-tag gels

Phosphorylation of TAB1 was detected using Phos-tag gels (Wako Pure Chemical Industries) containing 100 µM Phos-Tag acrylamide and 100 uM MnCl_2_ according to the manufacturer’s instructions.

### Statistical Analysis

Data were analyzed and assessed for outliers and significance using Prism software (version 10; GraphPad software, La Jolla, CA). Significance was determined by using a one-way analysis of variance. Data visualization was accomplished by using the graphing feature. Comparisons of statistical significance were determined through either one-way or 2-way analysis of variance (ANOVA) as indicated. Significance indicated with asterisks for designate *p* values as indicated in legends.

## RESULTS

### Atypical P38 Suppression in the Tab1^KI^ Mice Blocks Retinal Damage

Previous studies have shown that mutating four key residues in the Tab1 c-tail in the Tab1^KI^ mice selectively blocks Tab1’s interaction with p38α and subsequent atypical p38 activation during ischemia-reperfusion injury (**Fig. 1A**)[29, 43]. To explore the impact of Tab1-dependent atypical p38 on retinal vascular damage, we used an oxygen-induced retinopathy (OIR) model (**Fig. 1B**). Briefly, pups at postnatal day 7 (pn7) were exposed to 70% oxygen (hyperoxia) for 5 days, inducing blood vessel regression (vaso-obliteration). At pn12, the mice were returned to room air (normoxia) for 5 days. The prior blood vessel regression causes a rapid drop in retinal oxygen due to reduced blood flow. The resulting hypoxic shock enhances HIF1α signaling, inducing VEGF expression, vessel damage, hyperproliferation, and the formation of pathological neovascular tufts. Retinas were collected at pn17 and compared to pn17 room air (RA) control mice. Retinal vascular complexity was first quantified from isolectin B4 (IB4) staining of retinal flat mounts using a Sholl mask analysis [35, 38]. RA wt and Tab1^KI^ mice displayed comparable vascular complexity, with no statistical differences (**Fig. 1Ci & ii and E, shaded light red and blue lines)**. In contrast, OIR wt and Tab1^KI^ mice showed reduced central retinal vascular complexity compared to RA mice (**Fig. 1Di & ii, E compare shaded lines to solid lines, see appendix 1 for the complete data set and significance analysis)**. However, OIR wt mice had significantly reduced vascular complexity compared to the Tab1^KI^ mice, indicating greater blood vessel loss (**Fig. 1Di versus Dii, and E**). We next quantified vessel loss and neovascular tufting using NIH ImageJ [37]. Wt mice showed characteristic vaso-obliterative damage (dashed line, **Fig. 1Fi**) and enhanced vascular proliferation (white spots, **Fig. 1Fi**). Both vaso-obliteration and neovascularization were reduced in the Tab1^KI^ mice (**Fig. 1Fii**). Vaso-obliteration decreased significantly from 15.7% +/- 3.7 in wt mice to 5.9%+/- 3.6 in Tab1^KI^ mice (**Fig. 1F, dashed line; G**). OIR wt mice also had significant levels (8.5% +/- 2.7 SD) of neovascular tufting (white spots, **Fig. 1Fi; H**), which were reduced to 5.4% +/- 2.3 SD in OIR Tab1^KI^ retinae (white spots, **Fig. 1Fii; H**). The size of neovascular tufts was additionally significantly reduced, from 3511.7 +/- 1311.6 to 1370.9 +/- 445.6 SD (**Fig. 1I**). Measurement of the blood vessels showed that Tab1^KI^ retinae had increased vascular length and branch points, but no significant difference in the vascular endpoints (tips) (**Fig. 1J-L**).

**Figure 1.**
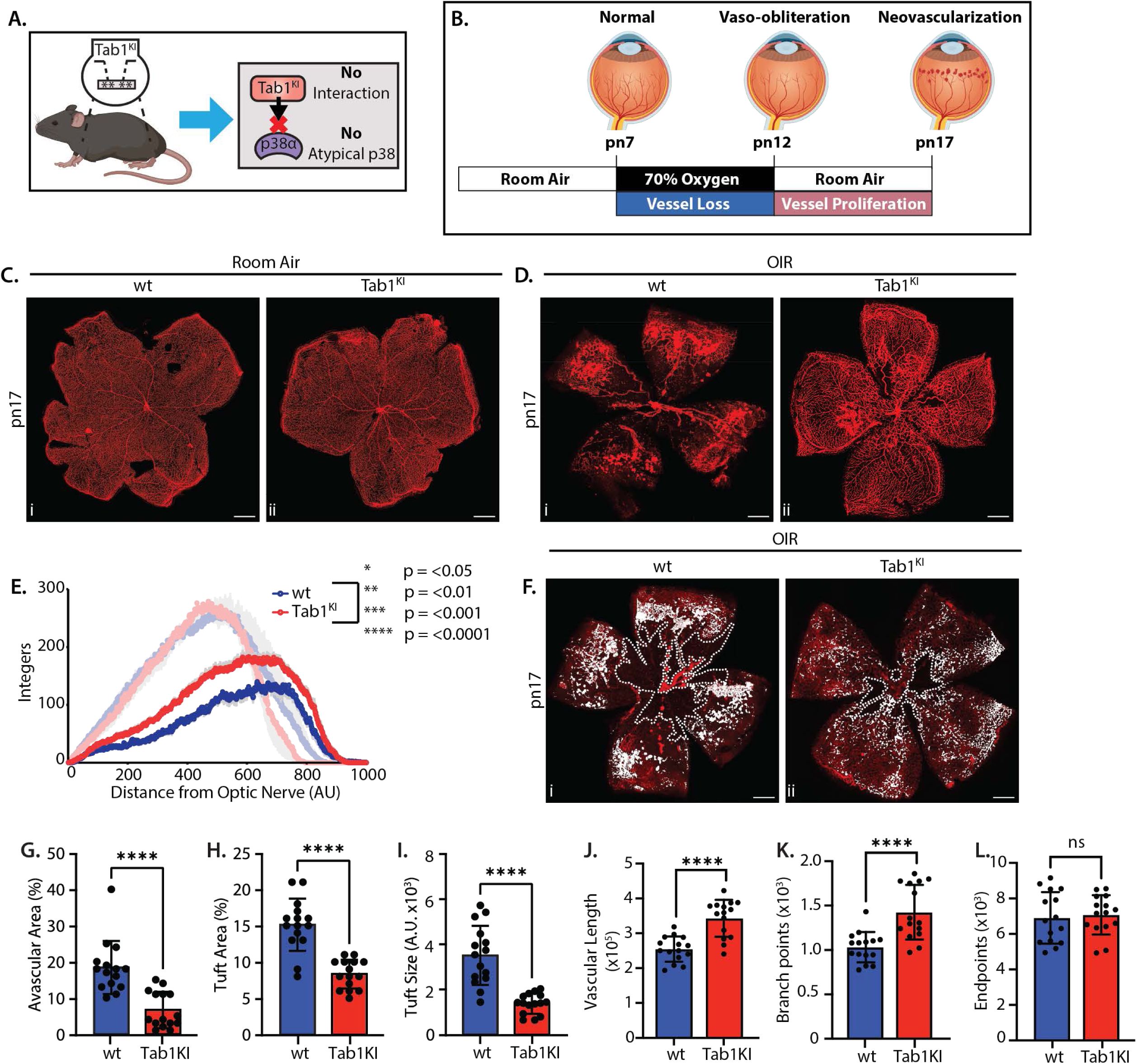
Vascular density and development in wt versus Tab1^KI^ mice after OIR. **(A)** Schematic model of Tab1^KI^ mice showing the four-point mutation blocking Tab1 interaction and atypical p38 signaling. **(B)** Timeline of OIR injury to mice from pn0 to pn17. **(C)** Retinal flat mounts of room air wt (i) and Tab1^KI^ (ii) mice with vasculature stained by Isolectin B4 (red). **(D)** Retinal flat mounts of OIR wt (i) and Tab1^KI^ (ii) mice with vasculature stained by Isolectin-B4 (red). **(E)** Sholl analysis graph of blood vessel density in OIR wt and Tab1^KI^ mice at radii of 1-1000 pixels from optic nerve (OIR n=6, RA n=4). Room air samples are underlaid for comparison. Significant values are displayed in **Supp. Table 1. (F)** Retinal flat mount of pn17 wt (i) and Tab1^KI^ (ii) mice showing avascular area (dashed outline) and neovascular tufts (white dots). Scale bar = 500 μm. **(G-L)** Quantification of the **(G)** avascular area, **(H)** tuft area, **(I)** tuft size, **(J)** vascular length, **(K)** branch points, and **(L)** endpoints in wt and Tab1^KI^ mice, each retina marked as a single data point. n = >13, were analyzed by t-test (****, *p* < 0.0001).

### Tab1^KI^ limits vascular tufts and disruption of the retinal ganglion cell layer

OIR induces robust disruption of the retinal vascular networks. To explore these changes in more detail, we first compared sectioned pn17 eyes from RA and OIR in both wt and Tab1^KI^ mice. RA wt and Tab1^KI^ mice showed no differences (**Fig. 2A, B**), but vascular tufting was significantly increased in wt OIR-treated mice (8.3 +/- 2.2 tufts per whole retinal section). This was reduced in Tab1^KI^ mice (3.3 +/- 1 tufts per section) (**Fig. 2C, D, E**). To verify vascular growth within the tufts, we stained tissue sections with the blood vessel marker IB4 (**Fig. 2 F)**. IB4 labelling confirmed normal blood vessel growth in the superficial, intermediate, and deep layers in both RA samples and OIR Tab1^KI^ retinae. In contrast, robust IB4 tuft staining was detected in OIR wt retinae. H&E staining also revealed that the retinal ganglion cell (RGC) layer was disrupted by vascular tufts in wt OIR mice and displayed additional nuclear stacking (**Fig. 2C**). To explore this further, we stained for RNA-binding protein with multiple splicing (RBPMS), an RGC marker, and CD31, a vascular endothelial marker. In both wt and Tab1^KI^ RA retinae, RBPMS clearly labeled a single well-defined RGC layer, and CD31 displayed the expected endothelial cell distribution (**Fig. 3Ai & ii**). In OIR wt retinae, RBPMS labelling demonstrated that the RGCs were embedded within the CD31-labelled vascular tufts (**Fig. 3Aiii**). In contrast, Tab1^KI^ retinae retained an even RGC distribution, similar to the H&E findings (**Fig. 3Aiv and 2D**).

**Figure 2.**
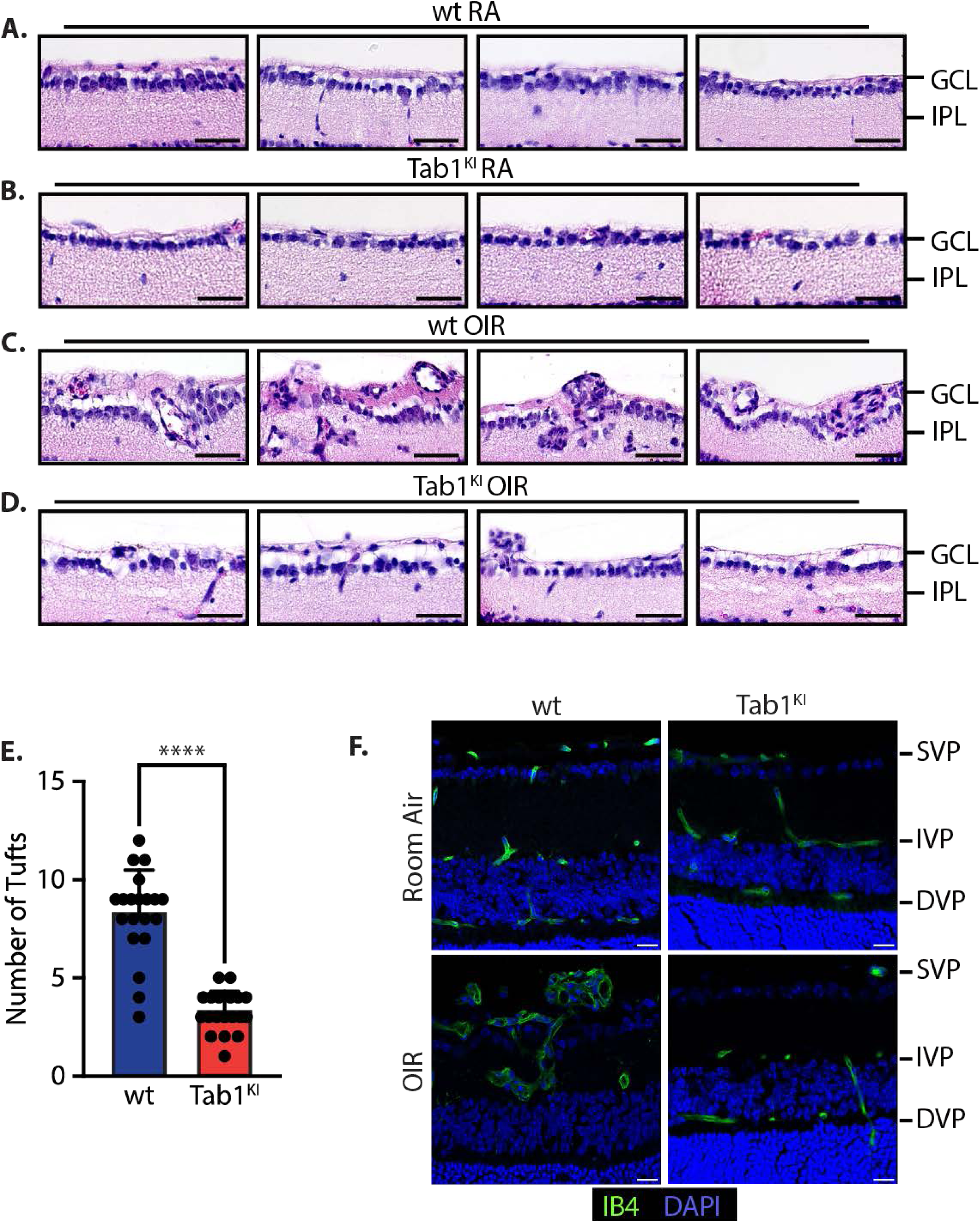
Histological and glial changes of wt versus Tab1^KI^ mice after OIR. **(A-D)** Hematoxylin and Eosin staining of all retinal layers in **(A)** room air wt mice, **(B)** room air Tab^KI^ mice, **(C)** OIR wt mice, and **(D)** OIR Tab1^KI^ mice. **(E)** Bar graph representing number of tufts (wt n=19, KI n=18) in retinal ganglion cell layer in OIR wt and Tab^KI^ mice. Each retinae image marked as a single data point. n = >13, were analyzed by t-test (****, *p* < 0.0001). **(F)** Representative immunofluorescence images of the retinal ganglion cell layer in room air wt and Tab1^KI^ mice and OIR wt and Tab1^KI^ mice stained with Isolectin-B4 (green) and DAPI (blue). Scale bar = 20 μm.

**Figure 3.**
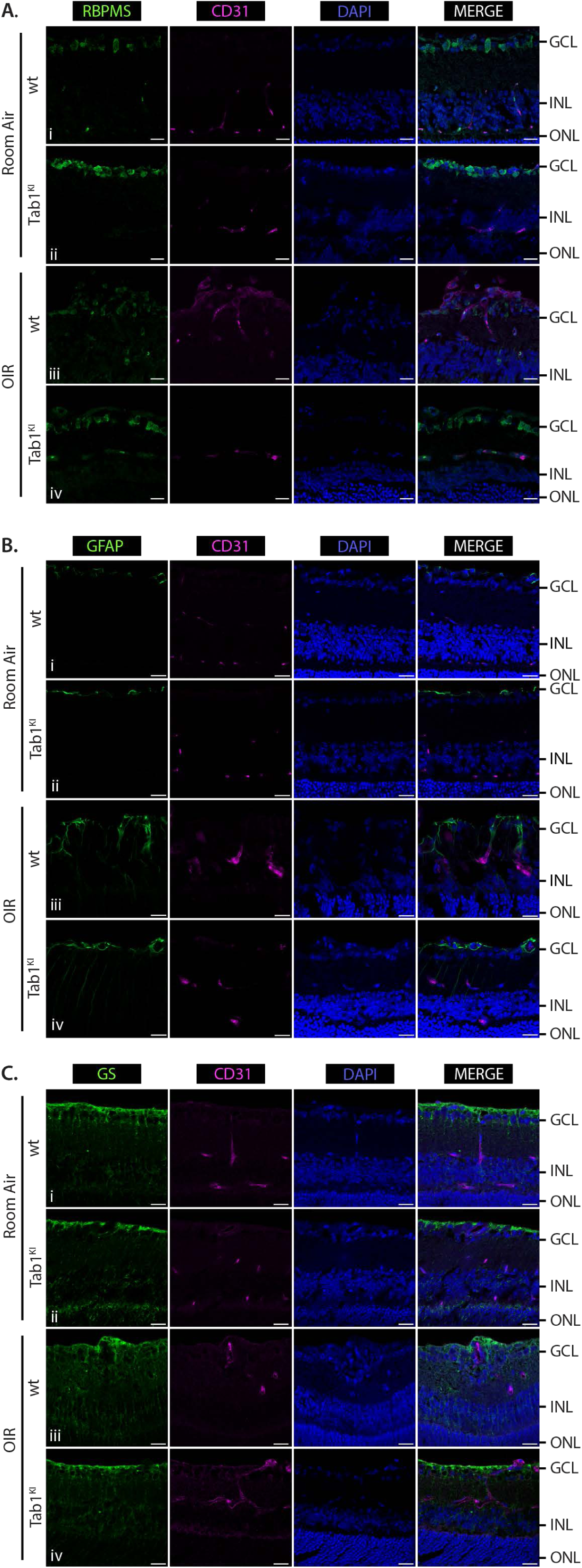
Expression of retinal ganglion and glial cells in wt and Tab1^KI^ mice after room air conditions and after OIR. Representative immunofluorescence images of retinal layers in room air and OIR wt and Tab1^KI^ **(A)** Immunofluorescence images of retinal layers in room air and OIR wt and Tab1^KI^ stained with RBPMS (green), CD31 (magenta), and DAPI (blue). **(B)** stained with GFAP (green), CD31 (magenta), and DAPI (blue). **(C)** stained with GS (green), CD31 (magenta), and DAPI (blue). Scale bar = 20 μm, n = >5.

Astrocytes located within the RGC layer facilitate vascular growth[44]. We therefore explored their location by labelling samples with glial fibrillary acidic protein (GFAP). In the RA retinae, GFAP-labelled astrocytes were confined to within or just above the RGC layer (**Fig. 3Bi & ii)**. In OIR retinae, GFAP staining was enhanced, indicating reactive gliosis as previously reported in astrocytes/Müller cells (**Fig. 3Biii & iv)**[45, 46]. Despite the relative protection from tuft formation in Tab1^KI^ retinae, GFAP labeling showed enhanced astrocytic projections (**Fig. 3Biv**). In wt retinae, these projections were overlapping with or immediately adjacent to enlarged CD31-positive vasculature (**Fig. 3Biii**). In contrast, in Tab1^KI^ retinae, astrocyte projections reached the inner nuclear layer but were not observed to be associated with CD31-positive blood vessels (**Fig. 3Biv**). Müller cells were also labelled with glutamate synthase (GS), consistent with the above GFAP observations, RA wt and Tab1^KI^ were nearly indistinguishable (**Fig. 3Ci and ii**). Besides a consistent rearrangement of the GS label, representing an exclusion from the vascular tufts in the wt OIR retinae, no robust variances were noted between the wt and Tab1^KI^ retinae after OIR (**Fig. 3Ciii and iv**).

Vascular tortuosity, used as an additional marker for retinal stress during OIR, refers to blood vessels with a curved or twisted morphology [47]. Comparing RA wt and Tab1^KI^ retinae, major arteries and veins showed similar linearity (**Fig. 4A, B and E**). In OIR wt retinae, retinal stress significantly reduced vessel linearity to ∼ 75% and increased vascular tortuosity (**Fig. 4C, E**). Despite reduced vascular tufting and vaso-obliteration in Tab1^KI^ retinae, tortuosity increases similarly, demonstrating an equivalent reduction in linearity (**Fig. 4D, E**).

**Figure 4.**
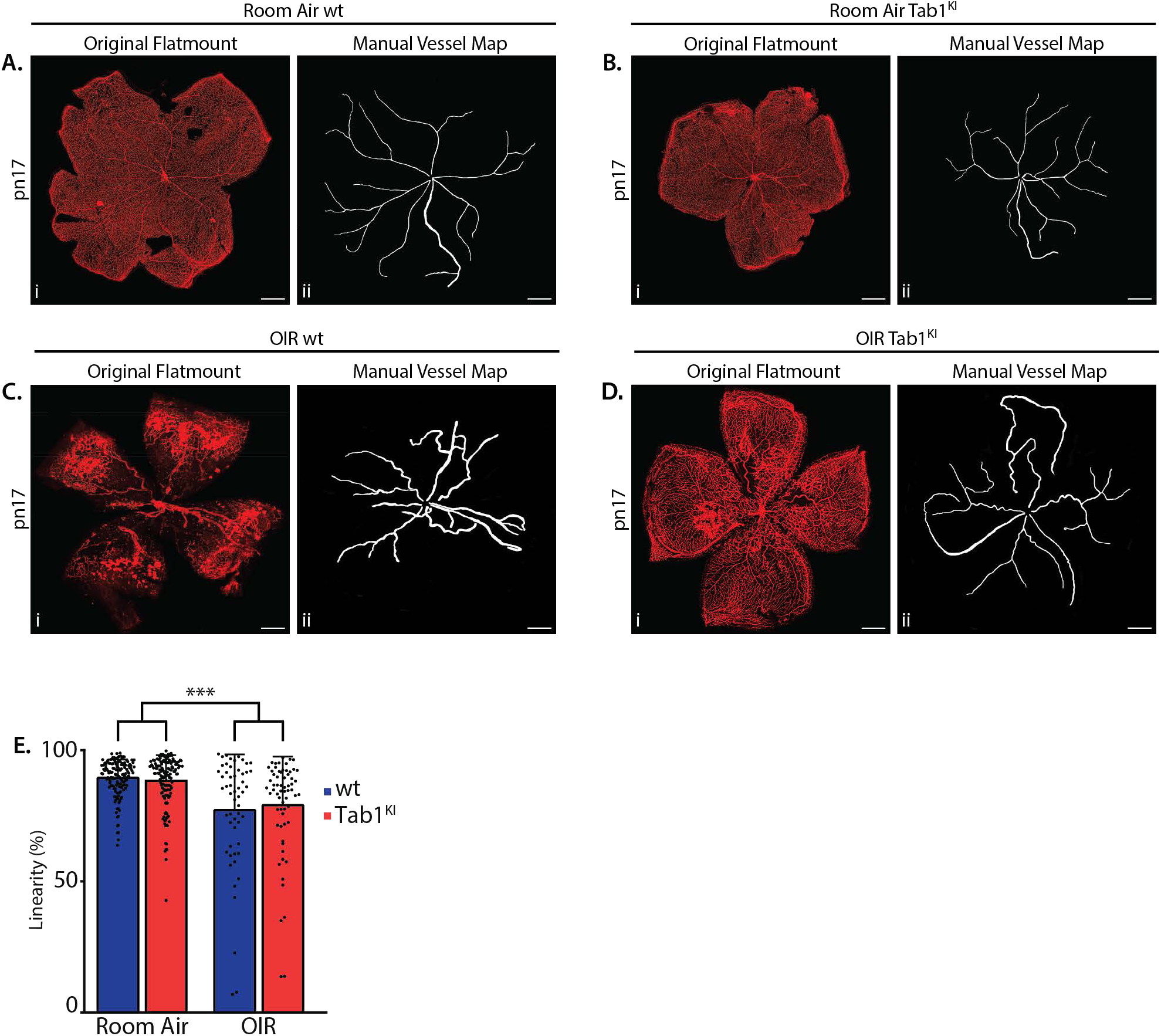
Vascular tortuosity in wt versus Tab1^KI^ mice in room air conditions and after OIR. **(A-D)** Retinal flat mounts of wt and Tab1^KI^ mice in room air and OIR conditions (i) and manual vessel maps (ii). Vasculature is stained using Isolectin B4 (red). **(E)** Bar graph representing % linearity of vessels (OIR wt n=42, OIR KI n=56, RA n=28, from n=5 retina each) in wt and Tab1^KI^ mice in room air and OIR conditions. Analyzed by T-Test (***, *p* < 0.001).

### Tab1 KI Confers Significant Transcriptional Changes in the OIR Retinae

We next compared the transcriptional profiles of wt and Tab1^KI^ retinae using RNAseq analysis of age and litter-matched retinae at pn17 under RA or OIR conditions (**Fig. 5A**). Replicate clusters were visualized with Uniform Manifold Approximation and Projection (UMAP) analysis, RA wt and Tab1^KI^ retinae displayed similar gene set complexity with overlapping characteristics. However, after OIR, the UMAP clusters showed distinct separation between their gene sets, demonstrating a divergence in transcriptional landscapes (**Fig. 5B**).

**Figure 5.**
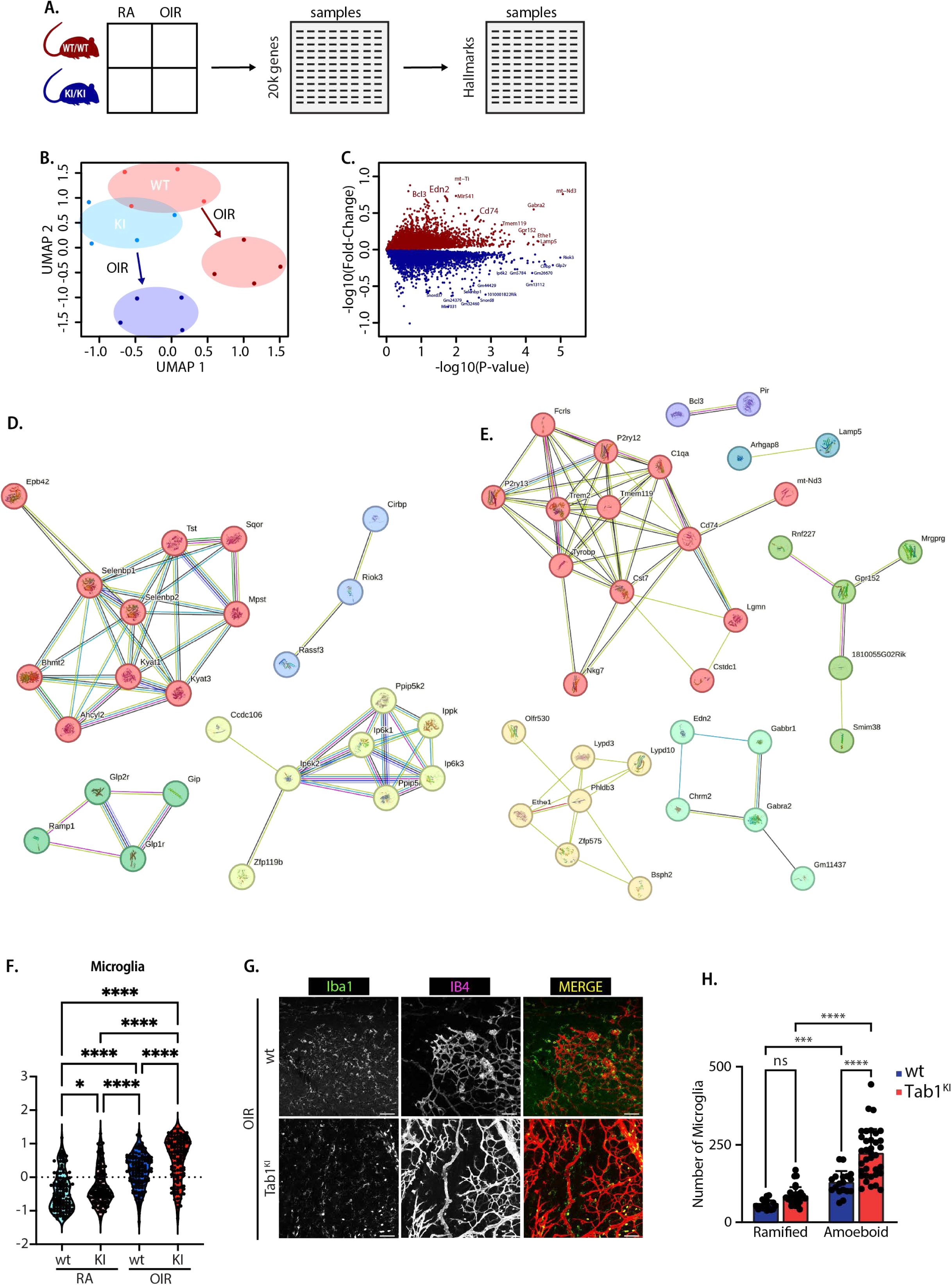
Tab1 KI Confers Significant Transcriptional Changes in the OIR Retinae. RNAseq analysis of wt or Tab1^KI^ Retinae, from pn17 RA or pn17 OIR mice **(A)** Schematic of the data organization and analysis. **(B)** UMAP comparative analysis of data sets, for wt (pink/red), Tab1^KI^ (blue/purple, KI) from room air (RA) and OIR retinae. **(C)** Volcano plot of raw differential expression showing max var genes: stress (mitochondrial genes) and immune genes (e.g. CD74). **(D & E)** String pathway analysis of the top genes from C, **(D** decreasing genes **& E** increasing genes**). (F)** Comparison of GSEA microglial marker gene expression, **(G)** Representative immunofluorescence images of retinal microglia (green) and IB4, vasculature (red). Scale bar = 20 μm. **(H)** Quantification of ramified and amoeboid microglia (wt n=18, KI n=36) in wt and Tab1^KI^ mice in OIR retinae.

Comparing the top differentially expressed genes (DEGs) between the OIR retinae, 399 genes were significantly altered (**Fig. 5C**). Pathway analysis of these top DEG clusters using STRING suggested a reduction in inositol phosphate biosynthesis, synthesis of pyrophosphates, Class B/2 Secretin family receptors, and sulfur metabolism in the Tab1^KI^ retinae (**Fig. 5D**). Conversely, genes associated with G protein-coupled neurotransmitter receptor activity, protein trafficking to the lysosome, and microglial activity were enhanced in the Tab1^KI^ retinae (**Fig. 5E**). To support this analysis, we compared the z-score normalized relative expression of microglial marker genes using gene sets derived from the Molecular Signature Database (MSigDB) (PMID: 26771021). Both wt and Tab1^KI^ RA retinae displayed comparable microglial marker expression, with a significant increase observed in OIR retinae of both genotypes. Notably, microglial marker expression increased significantly more in Tab1^KI^ retinae than wt retinae (**Fig. 5F**). These mRNA associations were evaluated at the protein level via co-labelling of the OIR retinal flat mounts from wt and Tab1^KI^ mice with IB4 (blood vessels, red) and allograft inflammatory factor 1, Iba1 (Microglia, green), we detected no significant difference in ramified microglia, but a significant increase in activated ameboid Microglia (**Fig. 5G and H**).

Examination of the MSigDB Hallmark transcriptional pathways signatures revealed significant differences between the RA and OIR samples and wt and Tab1^KI^ retinae (**Fig. 6A**). Rather than showing a profile similar to RA, the Tab1^KI^ OIR retinae presented a distinct pattern of pathway changes relative to the wt OIR samples. Because p38 is a proinflammatory kinase, we predicted that the reduction in vaso-obliteration and vascular tufting would coincide with a decrease in OIR-induced inflammation. However, Tab1^KI^ retinae displayed an enrichment of inflammatory signaling pathways, including IL6/STAT3, IFN-γ, IFN-α, TNFα, and NF-kB pathway genes (**Fig. 6A, B**).

**Figure 6.**
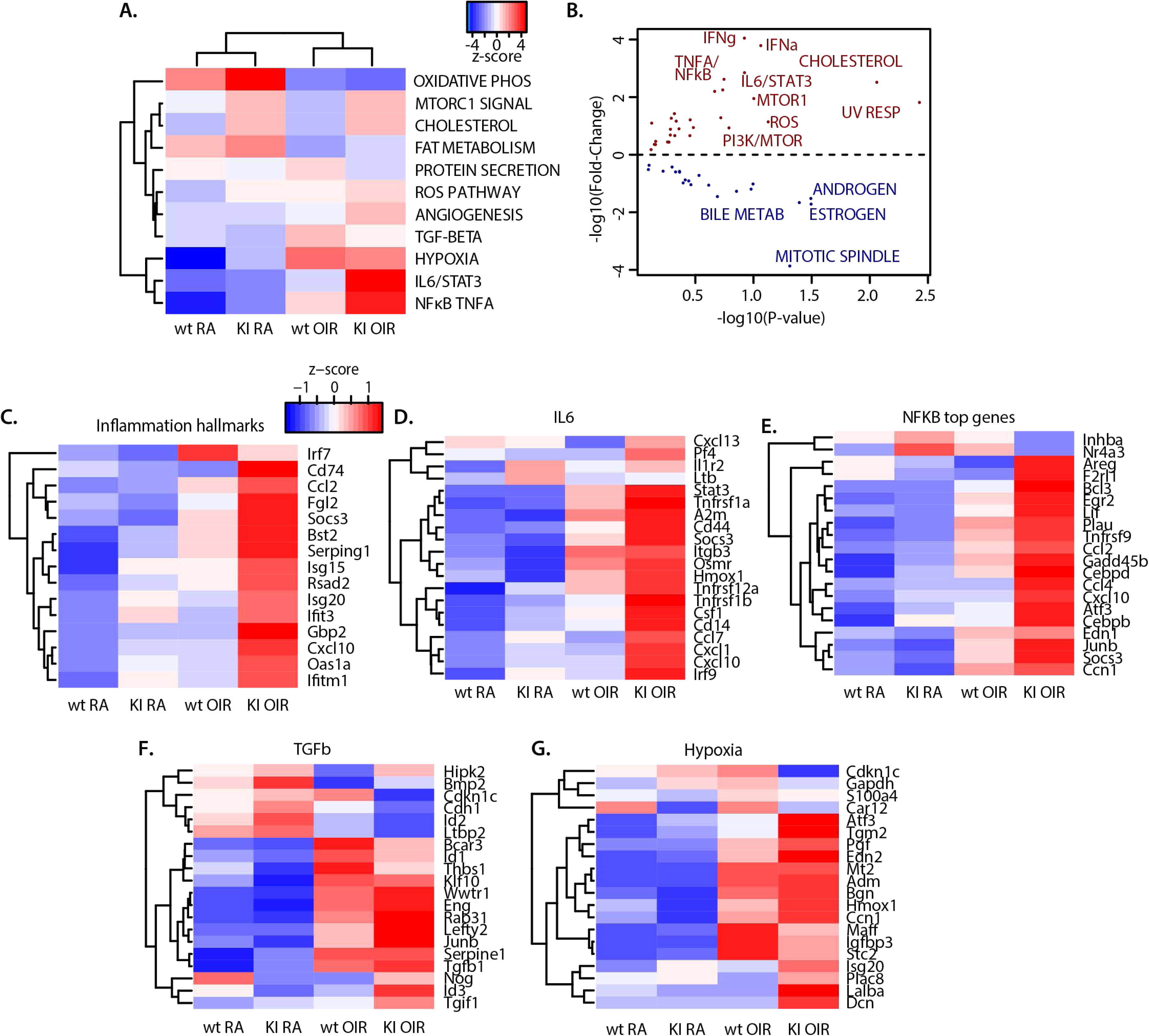
Analysis of GSEA hallmark pathway gene sets. RNAseq analysis of wt or Tab1^KI^ Retinae, from pn17 RA or pn17 OIR mice **(A)** Selected examples of the GSEA hallmark gene sets. **(B)** Volcano plot of pathway heatmaps representing fold change relative to significance. **(C-G)** Top 20 significantly increased genes for; **(C)** Inflammatory hallmarks **(D)** IL6 pathway **(E)** NFκΒ pathway **(F)** TGF-Β pathway **(G)** Hypoxia

Further examination of inflammatory genes revealed increased inflammation in the Tab1^KI^ retina samples after OIR (**Fig. 6C**). A closer examination of the top 20 genes associated with IL-6 and NF-κB signaling showed increases in both wt and Tab1^KI^ retinae after OIR, with Tab1^KI^ retinae exhibiting a robust increase (**Fig. 6D, E**). Similarly, markers for hypoxia and TGFβ signaling pathways were elevated in both wt and Tab1^KI^ retinae after OIR (**Fig. 6F, G**).

### P38 activity is significantly altered during OIR in the absence of Tab1-dependent p38 signaling

To explore changes in published p38 activation transcriptional-signatures between wt and Tab1^KI^ retinae in RA and OIR retinae, we initially compared a comprehensive set of 12 p38 gene signatures from the molecular signature database (MsigDB) (**Fig. 7A**)[48]. Out of the 12 representative data sets, there are distinct differences in p38-activated genes between wt and Tab1^KI^ retinae, both in RA and the OIR samples. Notably, rather than seeing a global loss of p38 signatures in the OIR Tab1^KI^ retinae, the apparent p38 markers are increased in both RA and OIR (**Fig. 7B**). However, p38-sig2 and p38-sig3 selectively increased in the wt OIR samples without a corresponding increase in the Tab1^KI^ OIR retinae. Comparing the OIR samples, out of the 889 p38-associated genes[48], 574 were specifically increased in the Tab1^KI^ retinae, 248 in wt retinae, and 67 in both datasets (**Fig. 7C**). Using the Biocarta p38 signaling pathway geneset, the z-scores of p38-associated genes displayed a significant increase between RA wt and Tab1^KI^ retinae. Furthermore, p38-associated genes were significantly increased in OIR relative to RA retinae for both wt and Tab1^KI^ mice (**Fig. 7D**).

**Figure 7.**
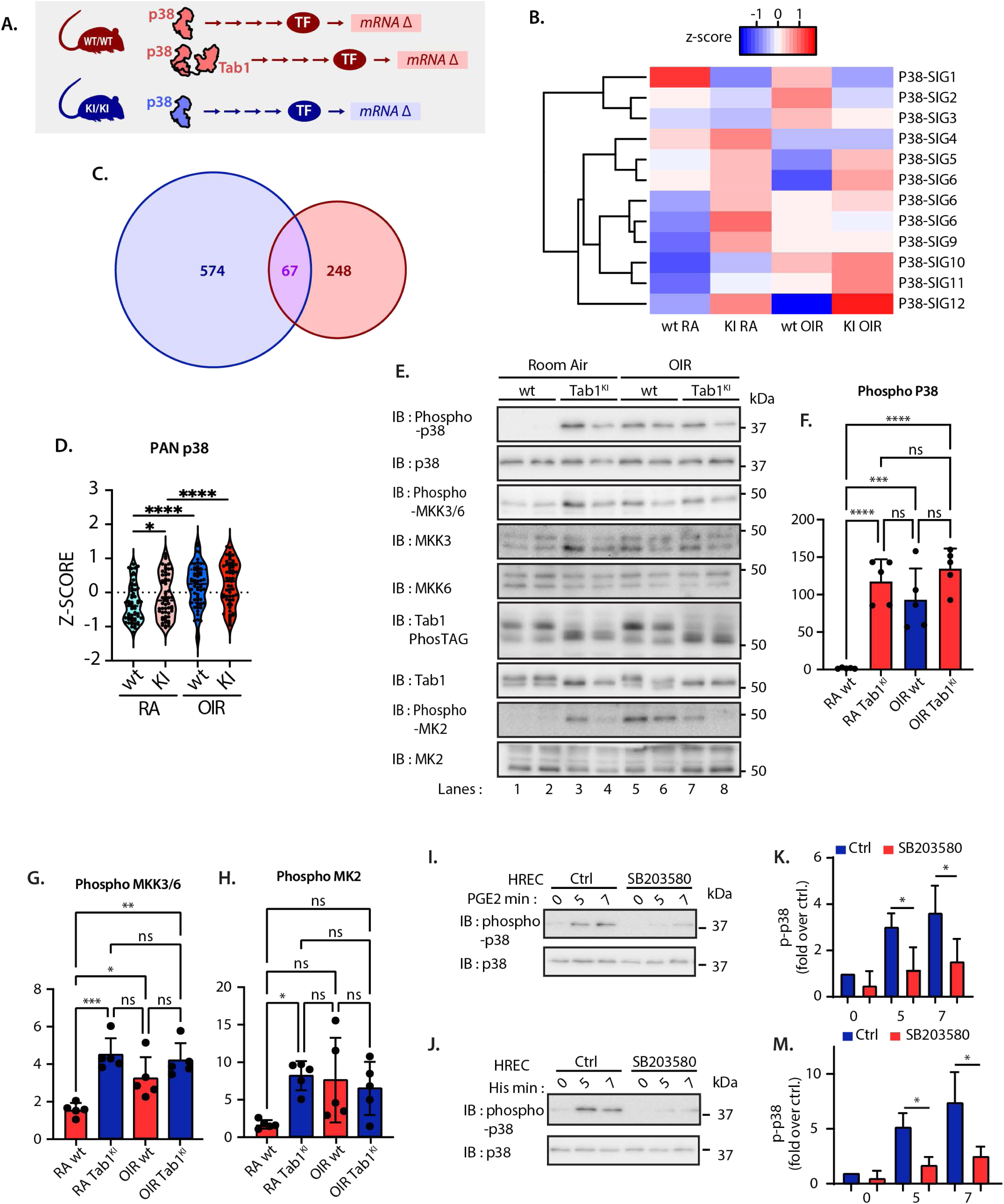
P38 activity in wt and Tab1^KI^ Retinae. RNAseq analysis of wt or Tab1^KI^ Retinae, from pn17 RA or pn17 OIR mice **(A)** A schematic flowchart illustrating distinct data explored: TF’s and mRNA-signatures associated with wt or tab1^KI^ retinae. **(B)** Comparison of GSEA published p38-gene expression barcodes. **(C)** Venn diagram of overlapping genes between wt and Tab1^KI^ retinae. **(D)** Example Biocarta p38 signaling gene set, graphing of Z-score comparisons between RA, OIR, wt, and Tab1^KI^ retinae. **(E)** Immunoblotting of RA, OIR, wt and Tab1^KI^ retinae, anti-phospho-p38, anti-p38, anti-phospho-MKK3/6, anti-MKK3, anti-MKK6, anti-Tab1 PhosTAG, anti-Tab1, anti-phospho-MK2, anti-MK2 **(F-H)** quantification of immunoblots from retinae **(F)** Fold change of phospho-p38 over total p38 quantified, mean +/- StDev, (n=5). **(G)** Fold change of phospho-MKK3/6 over total MKK6 quantified, mean +/- StDev, (n=5). **(H)** Fold change of phospho-MK2 over total MK2 quantified, mean +/- StDev, (n=5). **(I/J)** Representative immunoblot of HREC cells and stimulated with 10 µM PGE_2_ (I) or 1 µM histamine (J) for the indicated times in the presence of p38 inhibitor 10µM SB203580 or DMSO control. **(K, M)** Fold change of phospho-p38 over total p38 quantified from three independent repeats, for PGE_2_ (K) or histamine (M) (mean +/- StDev). All Immunoblots were analyzed by One-Way ANOVA (*, *p* < 0.05, **, *p* < 0.01, ***, *p* < 0.001, ****, *p* < 0.0001).

Contrary to our expectation, these data suggest that rather than blocking all p38 activity, the Tab1^KI^ retinae retain significant levels of p38 activity. Immunoblotting of wt and Tab1^KI^ retinae demonstrated that activation of p38 was significantly increased in RA Tab1^KI^ retinae, with no further increase after OIR, as indicated by detection of the dual (Thr180-Tyr182) phosphorylation of p38, (**Fig. 7E, compare lanes 1-2 with 3-4 and 7-8, quantified in 7F)**. Conversely, p38 phosphorylation was absent in wt retinae, but increased after OIR (**Fig. 7E, compare lanes 1-2 with 5-6, quantified in 7F)**. We next determined whether the observed increase in phospho-p38 was due to enhanced classical p38 signaling via MKK3/6. Indeed, phospho-MKK3/6 was significantly increased in RA Tab1^KI^ retinae, and as seen above for phospho-p38, OIR induced phospho-MKK3/6 in wt OIR with no additional increase in the Tab1^KI^ retinae (**Fig. 7E, compare lanes 1-2 with 5-6, quantified in 7G)**. Correspondingly, a well-established p38 target, MAPKAPK2, or MK2, was also found to be increased in the Tab1^KI^ samples, mirroring the MKK3/6 and p-p38 signals (**Fig. 7E, compare lanes 1-2 with 5-6, quantified in 7H)**. As previously shown, Tab1 is phosphorylated at multiple sites [10, 49, 50], and disruption of the Tab1-p38 interaction in the Tab1^KI^ mouse blocks p38-dependent Tab1 phosphorylation [29]. Tab1 migrated as a single band in the Tab1^KI^ retinae, but as two bands in the wt retinae (**Fig. 7E, compare lanes 1-2 with 3-4)**. As previously shown, Phos-TAG™ labelling of Tab1 (Phos-TAG^TM^ is a compound that selectively binds phosphorylated proteins in gels) confirmed that the wt retinae have a higher molecular weight phosphorylated Tab1 [32], which is absent in the Tab1^KI^ retinae (**Fig. 7E, compare lanes 1-2 with 3-4)**; of note, no additional Tab1 phosphorylation was observed after OIR. These data establish that while Tab1^KI^ perturbs the p38-mediated Tab1-phosphorylation and p38 autoactivation, it enhances classical, MKK3/6-dependent signaling in the retina. These data demonstrate that Tab1^KI^ does not suppress all p38 activity. But instead, the retina compensates for the loss of Tab1-dependent p38 activation by upregulating Mkk3/6 activity and “classical” p38 responses.

Prior studies have established atypical p38 signaling in primary human endothelial cells, using the proinflammatory and angiogenic histamine or prostaglandin E2 (PGE_2_)[31]. PGE_2_ activation has also been linked to pathological retinal neovascularization [31, 32, 51]. We next wanted to confirm that atypical p38 activity is active in primary human retinal endothelial cells (HREC). The acute stimulation of HREC with either PGE2 or histamine induced robust p38 activation (**Fig. 7I, J lanes 1-3)**. The p38-selective ATP-competitive inhibitor SB203580 blocks p38 kinase activity and prevents agonist-dependent p38 autophosphorylation/activation, which is critical in the Tab1-dependent atypical p38 activation pathway [31, 32]. Preincubation of HREC with 10µM SB203580 significantly suppressed both PGE_2_-and histamine-induced p38 autophosphorylation (**Fig. 7I, J, compare lanes 1-3 with 4-6, quantified in K and M)**. These data confirm that GPCR-induced atypical p38 is conserved in human retinal endothelial cells, emphasizing our findings in our OIR model.

### Atypical P38 Suppression Selectively Regulates Transcriptional Activity After OIR

As shown above, rather than suppressing all p38 activity, blocking the Tab1-dependent activation enhanced retinal p38 responses. Specifically, Tab1-dependent p38 activation selectively regulates a subset of inflammatory signaling pathways. In Fig. 7B & C, we show that p38-associated genes are substantially dysregulated after OIR. A closer examination of all atypical-genes reveals that a large group of genes is selectively upregulated in wt OIR retinae relative to Tab1^KI^ retinae (**Fig. 8A, top cluster).** Focusing on the GSEA Biocarta p38 panel, a subset of genes displayed a slight increase in Tab1^KI^ RA retinae but did not increase after OIR, conversely, wt retinae exhibit a significant increase in these genes after OIR relative to both RA wt samples and Tab1^KI^ OIR retinae (Fig. 8B).

**Figure 8.**
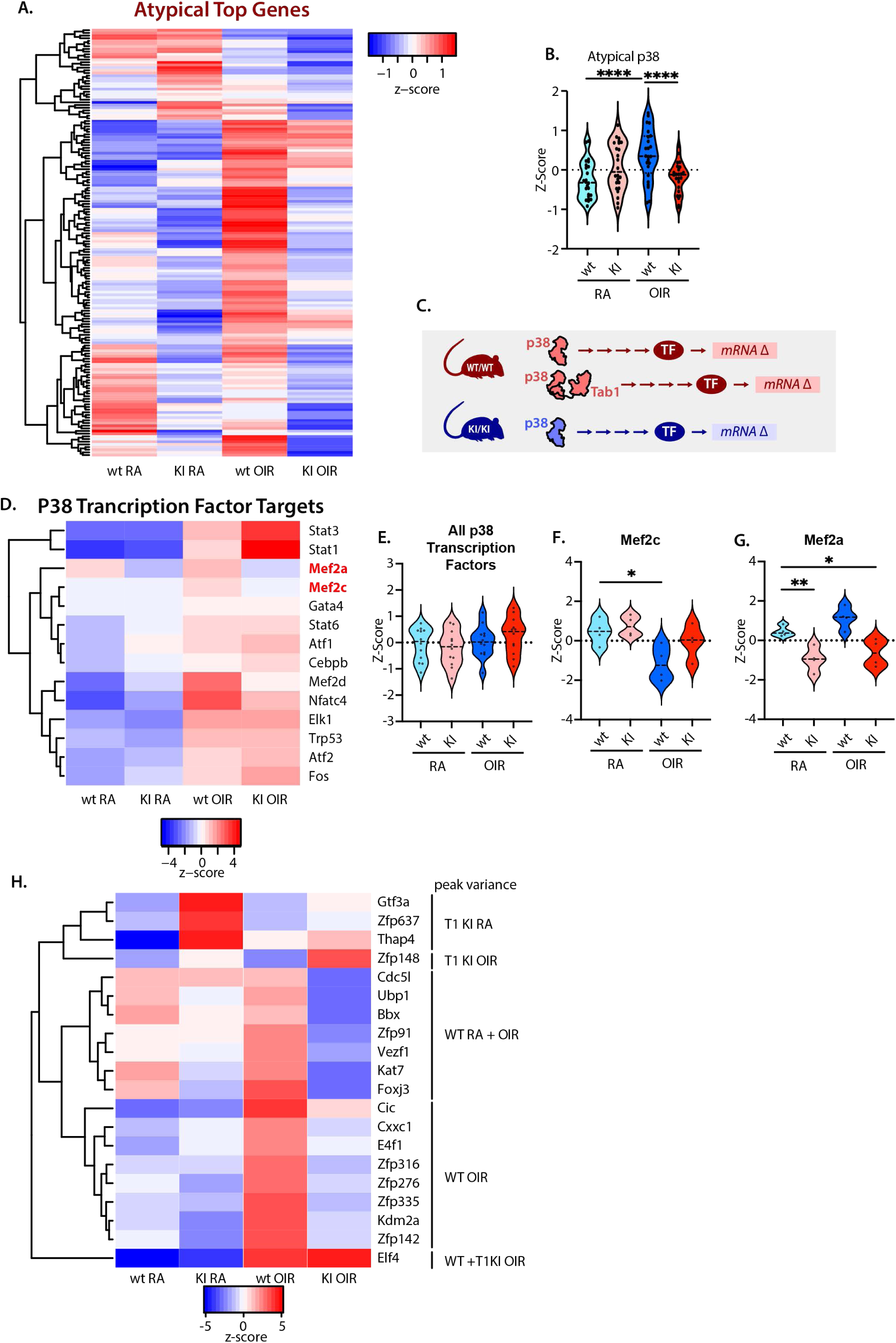
Atypical p38 suppression in the Tab1^KI^ OIR significantly changes p38-dependent transcriptional outcomes. RNAseq analysis of wt or Tab1^KI^ Retinae, from pn17 RA or pn17 OIR mice **(A)** Z-score heatmap for genes with higher expression in wt OIR than Tab1KI OIR retinae **(B)** Example genes from Biocarta p38 signaling gene set, graphing of Z-score comparisons between RA, OIR, wt, and Tab1^KI^ retinae, where there is an increase in wt OIR relative to RA, but downregulation in Tab1KI OIR retinae. **(C)** A schematic flowchart illustrating distinct data explored: TF’s and mRNA-signatures associated with p38 activation. **(D)** Z-score heatmap for activity of TF’s known to be downstream of p38 **(E)** Z-score, relative expression of p38-regulated TFs from **D**, **(F)** Z-score, relative expression of Mef2C, **(F)** Z-score, relative expression of Mef2A **(G)** Z-score heatmap for top TF’s activity for each experimental condition. Analyzed by One-Way ANOVA (*, *p* < 0.05, **, *p* < 0.001, ****, *p* < 0.0001).

The differential control of p38 target genes likely reflects regulation of transcription factors. We next examined the regulation of transcription factor signaling pathways (**Fig. 8C**). When comparing 13 p38-regulated transcription factors, we found no robust differences in the RA retinae. However, signaling was increased for eleven of these after OIR induction in both wt and Tab1^KI^ retinae (**Fig. 8D**). Notably, the signaling pathways for two key p38 transcription factors, Myocyte enhancer factor 2 A (Mef2A) and Mef2C, were suppressed (**Fig. 8D**). As a group, the mRNA profiles of the 13 transcription factors were not significantly different under any condition (**Fig. 8E**). Although Mef2C was reduced in the wt OIR retinae (**Fig. 8F**), and Mef2A was reduced in both the RA and OIR samples, this reduction was observed only in the Tab1^KI^ retinae (**Fig. 8G**). In addition to the known p38 transcription factors, we found that multiple additional transcription factor pathways displayed genotype-specific variances (**Fig. 8H**, with the sample showing the largest peak variance noted, i.e., the top 3 showing an increase in Tab1^KI^ RA mRNA gene expression), demonstrating broader impacts of atypical p38 beyond the factor pathways. These results indicate that Tab1-dependent atypical p38 signaling, in contrast to classical MKK3/6-driven activation, selectively shapes retinal transcriptional programs, providing a key mechanism for fine-tuning inflammatory and stress responses after vascular injury.

### Tab1KI OIR-Angiogenic Responses

As stated above, the Tab1^KI^ retinae displayed a significant reduction in OIR-induced retinal obliteration and neovascularization (**Fig. 1C-L**). We next examined the angiogenic pathways in the RNAseq samples. Wt and Tab1^KI^ retinae expressed comparable angiogenic markers under RA conditions, with the predicted significant increase after OIR in wt retinae. Contrary to our expectations, despite the reduced neovascular tufts, Tab1^KI^ retinae displayed a significant increase in angiogenic markers, above that seen in wt OIR retinae (**Fig. 9A**). Examining a panel of angiogenic factors and receptors, a similar trend for increased expression after OIR was observed (**Fig. 9B**). Notably, VEGFA was equivalently expressed and increased significantly after OIR in both the wt and Tab1^KI^ retinae (**Fig. 9C**). Furthermore, VEGF receptor 1, VEGFR1 (also known as Fms-related tyrosine kinase-1, Flt1), was significantly increased in wt OIR but not Tab1^KI^ retinae (**Fig. 9D**), whereas VEGFR2 (also known as kinase insert domain receptor, KDR), the main driver of blood vessel growth, was significantly increased only in Tab1^KI^ OIR retinae (**Fig. 9E**). Of note, IL19, previously shown to control OIR induced neovascularization, was increased only in the wt OIR sample, albeit not significantly (**Fig. 9B and F**) [52].

**Figure 9.**
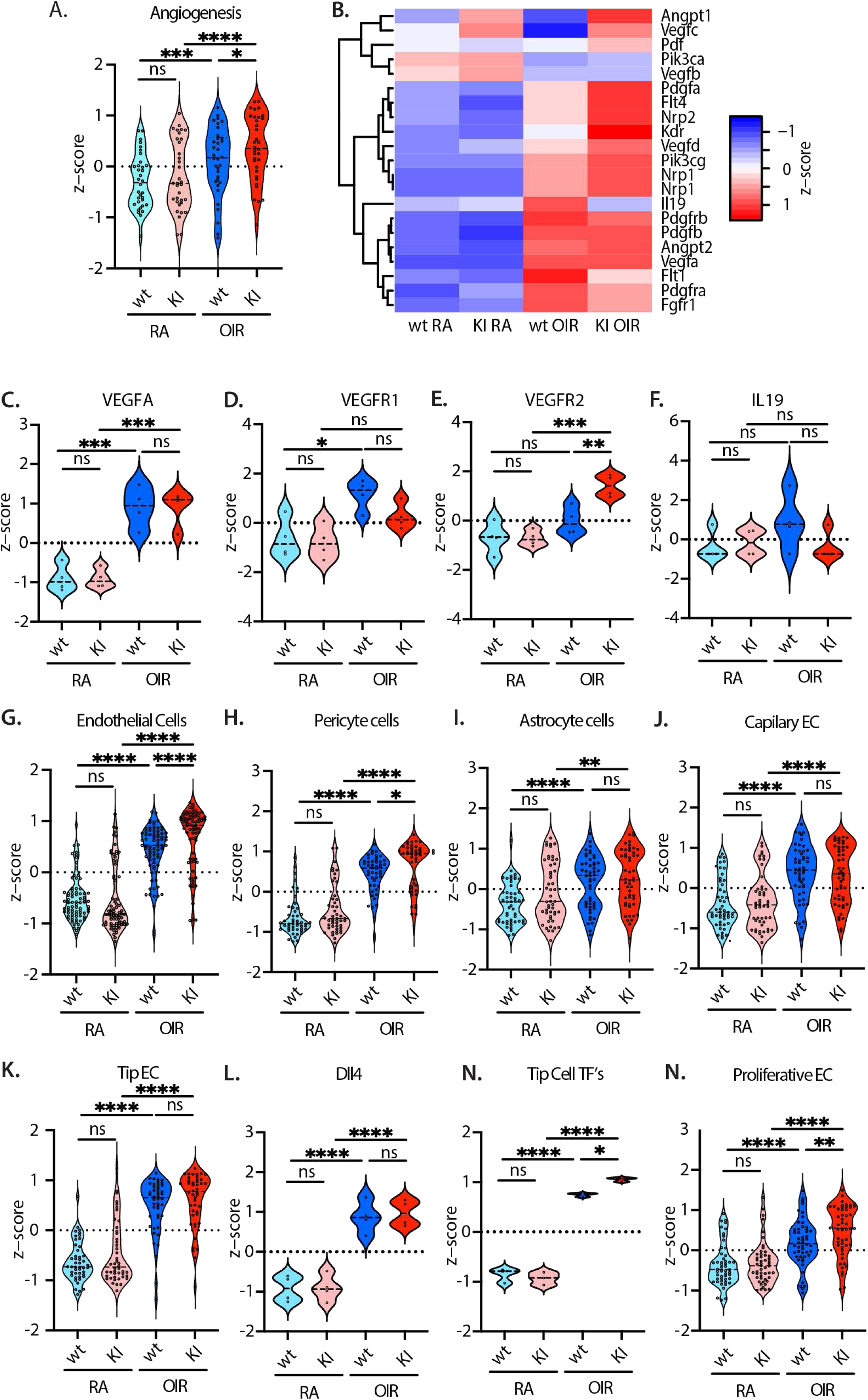
Regulation of Angiogenic Signaling in wt and Tab1KI Retinae. RNAseq analysis of wt or Tab1^KI^ Retinae, from pn17 RA or pn17 OIR mice **(A)** Z-score, relative expression of GSEA angiogenic markers. **(B)** Z-score heatmap for key genes in retinal angiogenesis. **(C)** Z-score, relative expression of VEGFA from **B. (D)** Z-score, relative expression of VEGFR1 from **B. (E)** Z-score, relative expression of VEGFR2 from **B. (F)** Z-score, relative expression of IL19 from **B. (G)** Z-score, relative expression of endothelial cell marker genes. **(H)** Z-score, relative expression of pericyte cell marker genes. **(I)** Z-score, relative expression of astrocyte cell marker genes. **(J)** Z-score, relative expression of capillary endothelial cell marker genes **(K)** Z-score, relative expression of tip endothelial cell marker genes **(L)** Z-score, relative expression of Dll4. **(M)** Z-score, relative expression of tip cell transcription factors, Elk4, Est1 and Hlx. **(N)** Z-score, relative expression of proliferative endothelial cell marker genes. Analyzed by One-Way ANOVA (*, *p* < 0.05, **, *p* < 0.001, ****, *p* < 0.0001).

As shown above, Tab1^KI^ retinae display a protective phenotype after OIR, limiting tuft formation and vaso-obliteration (**Fig. 1C-L**). Consistent with the flat mount analysis, endothelial and pericyte marker genes were significantly increased after OIR. The Tab1^KI^ retinae also displayed an additional significant increase above that observed in wt OIR retinae (**Fig. 9G and H**). This correlates with the enhanced vascular networks in the Tab1^KI^ retinae. In contrast, the astrocyte markers were increased after OIR in both wt and Tab1^KI^ retinae (**Fig. 9I**). Interestingly, further analysis of the endothelial genes revealed that capillary endothelial cell and endothelial tip cell genes were equivalently increased in both wt and Tab1^KI^ OIR retinae (**Fig. 9J and K, respectively**). Whereas, proliferative endothelial cell markers were significantly increased after OIR, with the Tab1^KI^ retinae displaying an additional significant increase relative to wt retinae (**Fig. 9L**). Together, these data indicate that Tab1^KI^ retinae retain a protective angiogenic profile after OIR, by enhancing proliferative endothelial responses while limiting pathological tuft formation. These data reveal a critical mechanism by which Tab1-dependent signaling can selectively modulate vascular remodeling in the injured retina.

## Discussion

Defining the molecular signaling that drives neovascularization in both healthy and diseased blood vessels is essential if novel therapeutic targets are to be identified. One such underexplored pathway in retinal neovascularization is the Tab1-dependent atypical p38 activation. First identified by Ge *et al.* in 2002 [30], atypical p38 is distinct from the classical MKK3/6-dependent p38 activation, relying instead on the direct interaction of the c-terminal tail of Tab1 and p38 to induce autoactivation [29, 53]. Tab1-dependent atypical p38 is selectively active in multiple disease pathologies ranging from cardiac ischemic damage to viral replication [10].

In this study, we examined atypical p38 in the regulation of pathological retinal neovascular damage using the well-established OIR model. The proliferative retinopathies (PRs) are a leading cause of blindness, manifesting as enhanced, aberrant pathological endothelial proliferation. PR induces the formation of neovascular tufts that extend into the vitreous space, damaging the RGC layer and exacerbating edema. We discovered that genetic perturbation of atypical p38 in the Tab1^KI^ mice significantly blocked OIR-induced neovascular-tuft formation. This response was independent of retained OIR-dependent MKK3/6-driven p38 signaling. The current therapeutic candidates for suppressing pathological p38 activity are predominantly ATP-competitive small molecule inhibitors [10]. Because MKK3/6-dependent and Tab1-dependent signaling utilize the same p38α kinase, the current small molecule p38 inhibitors block all p38 activity, without discrimination for a specific pathway. Our data indicate a potential benefit from using selective atypical p38 inhibitors. At present, there are no atypical p38 selective inhibitors in clinical trials.

However, recent studies have identified potential small-molecule protein-protein interaction inhibitors [54], as well as p38 allosteric modulators that selectively block atypical p38 activation in vitro, which are promising candidates for future development [55].

Critically, while we show that suppression of atypical p38 signaling limited OIR-induced damage, there was no difference in the retinal vascular development. This is consistent with the prior studies, exploring ischemia reperfusion injury [29, 43] and acute lung injury (unpublished data), reporting no impact on cardiac or lung development (respectively).

The selective activation of atypical p38 during OIR suggests that the formation of the Tab1-p38 complex drives distinct signaling events that lead to pathological outcomes. One hypothesis is that the Tab1-p38 interaction alters access to and activation of transcription factors [56, 57]. Our data show that out of 14 transcription factors known to be activated by p38, only Mef2a and Mef2c gene targets are differentially decreased in the Tab1^KI^ OIR retinae. Mef2a and Mef2c have been linked to microglial activity and pathological neovascularization in OIR [58, 59]. As stated above, the Tab1^KI^ retinae exhibited significant increases in microglial marker genes and an increase in amoeboid microglia in the retina after OIR. Additionally, inflammatory markers, including IL6 signaling, were also increased in the OIR Tab1^KI^ retinae. These data are consistent with a recent study that describes Mef2c-dependent restraint of microglial activation. Hu. *et al.* 2025 established that suppression of Mef2c activity, either by genetic knockout or chemical inhibition, enhanced microglial activity and microglial-dependent inflammation, increasing interleukin (Il6), Il1α, Il1β, TNFα, Ccl2/Mcp-1, and Cxcl10 mRNA expression [59].

In addition to microglial activation, Mef2c regulates both developmental and pathological blood vessel growth [58, 60-62]. More specifically, Mef2c expression and activation can direct pathological retinal neovascularization [58]. Because of the critical developmental roles of Mef2c, the Xu *et al.* used endothelial-specific Mef2c ablation. In these mice, retinal vascular development was unimpaired. However, after OIR, vaso-obliteration was reduced and vascular tufting was suppressed, with Mef2c suppression inducing a switch from pathological vasculogenesis to physiological vascular regrowth. Atypical p38 suppression in the Tab1^KI^ mice mimicked these data, limiting vaso-obliteration and tuft formation.

VEGFA can induce Mef2c expression in a p38-dependent manner, and p38 has also been shown to activate Mef2c directly [60, 63]. It is important to note that Mef2c expression was not suppressed in the Tab1^KI^ retinae after OIR and that developmental vasculogenesis was not perturbed. This suggests that the atypical, Mef2c-driven OIR is pathologically selective, likely driven by a complex interplay of signaling pathways that trigger tuft formation.

Mef2c-dependent physiological regulation of vasculogenesis is still a subject of investigation [61-63]; however, Mef2c can control EC tip cell signaling, via the Notch ligand, delta-like ligand4 (Dll4), and cross-talk with the Mef2c-dependent transcription factors, Hlx, Elk3 and Est1 [61-64]. We show that, in addition to equivalent developmental vascular growth in the RA mice, after OIR, vascular endpoints (tips) and endothelial tip cell genes were comparable between wt and Tab1^KI^ retinae. More specifically, Dll4, Hlx, Elk3, and Est1 all increase in expression after OIR, in both mouse lines. Conversely, proliferative EC’s marker genes displayed a greater increase in the Tab1^KI^ retinae, which supports the hypothesis that atypical p38 suppression does not prevent physiological vascular regrowth and tip cell formation. Therefore, atypical p38 selectively enhances pathological tuft formation via Mef2c signaling. Critically, angiogenic markers and VEGFA expression are not suppressed in the Tab1^KI^ mice. One potential protective response ismediated by Angiopoietin-1, also known as Ang1. Ang-1 is a critical regulator of physiological angiogenic signaling, balancing vascular health and survival. Conversely, angiopoietin-2 (Ang2) is upregulated in pathological pathways that drive vascular destabilization and VEGFA-mediated pathogenesis [65, 66]. In our study, angiopoietin-1 (Ang1) was increased in Tab1^KI^ mice both in the control RA and OIR retinae relative to the wt mice. Conversely, Ang-2 was enhanced in both wt and Tab1^KI^ after OIR. Therefore, Ang-1 in the Tab1^KI^ mice could be supporting a protective environment for retinal repair after OIR.

In conclusion, we demonstrate that Tab1-dependent atypical p38 signaling is a significant driver of pathological responses during proliferative retinopathy, as observed in the OIR mouse model. Selective disruption of atypical p38 provides protection from retinal tuft formation, while supporting vascular repair. This is independent of retained MKK3/6-dependent 38 signaling in the retinae—selective perturbation of transcription factor activation and pathogenic vascular growth. Our Tab1^KI^ OIR model provides a rationale for the development of atypical p38 selective inhibitors, or allosteric modulators. Limiting Tab1-dependent p38 activity to enhance vascular repair, suppress vascular dysregulation, and support the current anti-VEGF therapeutics. Further studies are ongoing to explore the mechanisms of atypical p38-induced pathological angiogenesis.

## Data Availability Statement

All data and resources are available upon request.

## Author Contributions

Project conception and design (NJG, PSN, and HK), experiments performed by (NJG, HK, LS, AY, FL, SG, and SSG), Data analysis and interpretation by (NJG, PSN, HK, ED, LS, AY, and FL), drafting manuscript (NJG), editing, review, and approval of final manuscript (NJG, PSN, HK, ED, LS, AY, FL, SG, SSG).

## Supplemental material

**Supplemental Table 1.**
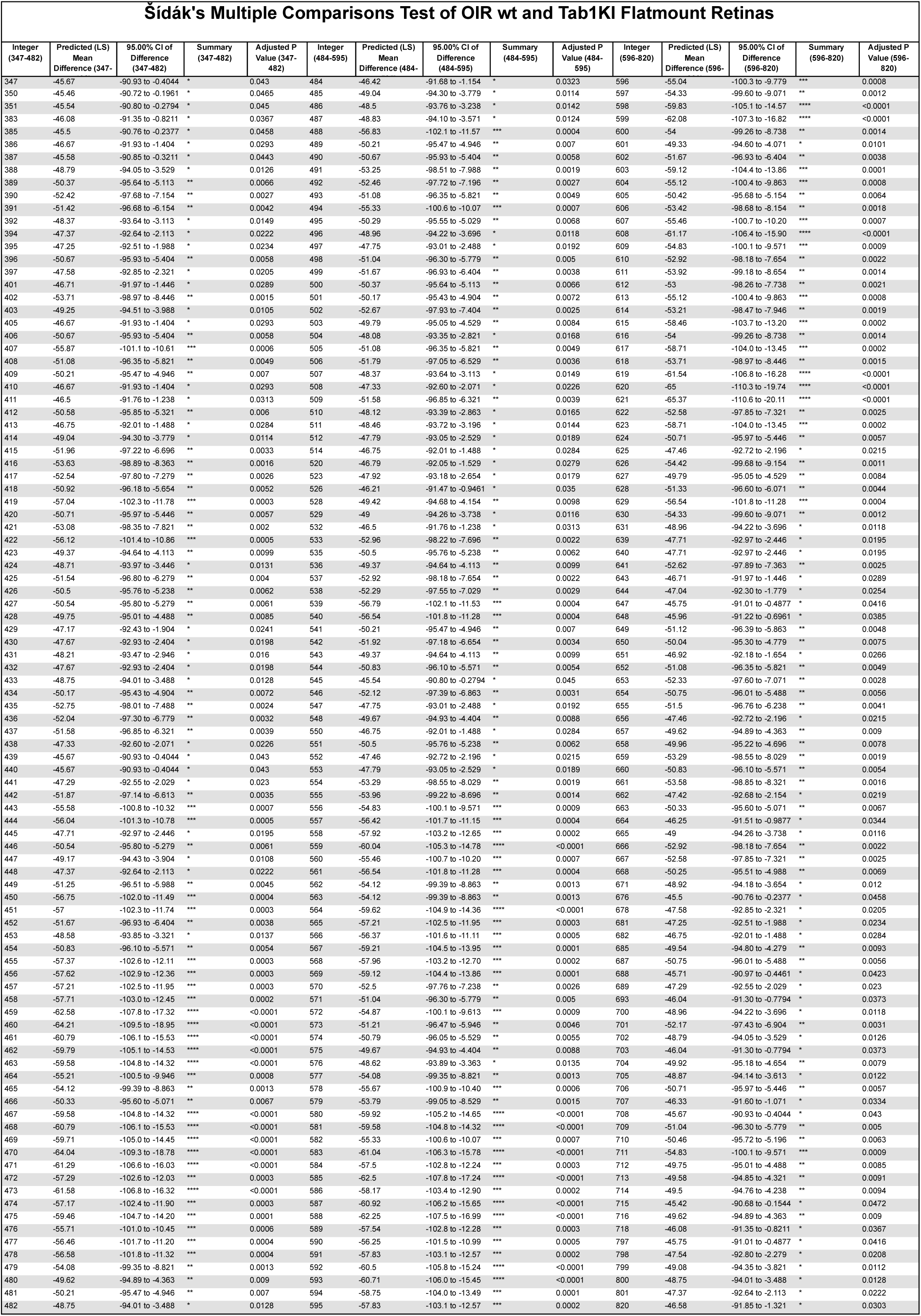
Statistical analysis from Sholl analysis of retinal flatmounts. Comparing wt and Tab1^KI^ retinae at pn17 after OIR, n=5 per genotype.

## Acknowledgments

We thank all members of the Grimseylab@UGA for their comments and advice.

## Footnotes

The following abbreviations were used;

AMD: Age-Related Macular Degeneration
DR: Diabetic Retinopathy
KI: Knock-In
IB4: Isolectin-B4
OIR: Oxygen-Induced Retinopathy
RA: Room Air
RBMPS: RNA-Binding Protein with Multiple Splicing
ROP: Retinopathy of Prematurity
Tab1^KI^: Tab1 Knock-In
VEGF: Vascular Endothelial Growth Factor
VEGFR: Vascular Endothelial Growth Factor Receptor
wt: Wild-Type
GFAP: Glial Fibrillary Acidic Protein
HREC: Human Retinal Endothelial Cell
RGC: Retinal Ganglion Cell

Others, not sure: NIH, OCT, (MKK3/6, Tab1, MK2, etc), EC, H&E, CD31, RNA seq stuff, SDS-PAGE, Phos-TAG, DAPI, GS,

